# Constraints and Opportunities for Detecting Land Surface Phenology in Drylands

**DOI:** 10.1101/2021.05.21.445173

**Authors:** Shawn D. Taylor, Dawn M. Browning, Ruben A. Baca, Feng Gao

**Affiliations:** US Department of Agriculture, Agricultural Research Service, Jornada Experimental Range, New Mexico State University, Las Cruces, New Mexico, 88003, USA; Oak Ridge Institute for Science and Education (ORISE), Oak Ridge, Tennessee, 37830, USA; US Department of Agriculture, Agricultural Research Service, Hydrology and Remote Sensing Laboratory, Beltsville, Maryland, 20705, USA

## Abstract

Land surface phenology (LSP) enables global scale tracking of ecosystem processes, but its utility is limited in drylands due to low vegetation cover and resulting low annual amplitudes of vegetation indices (VIs). Due to the importance of drylands for biodiversity, food security, and the carbon cycle it is necessary to understand limitations in measuring dryland dynamics. Here, using simulated data and multi-temporal unmanned aerial vehicle (UAV) imagery of a desert shrubland, we explore the feasibility of detecting LSP with respect to fractional vegetation cover, plant functional types, VI uncertainty, and two different detection algorithms. Using simulated data we found that plants with distinct VI signals, such as deciduous shrubs, can require up to 60% fractional cover to consistently detect LSP. Evergreen plants, with lower seasonal VI amplitude, require considerably higher cover and can have undetectable phenology even with 100% vegetation cover. Our evaluation of two algorithms showed that neither performed the best in all cases. Even with adequate cover, biases in phenological metrics can still exceed 20 days, and can never be 100% accurate due to VI uncertainty from shadows, sensor view angle, and atmospheric interference. We showed how high-resolution UAV imagery enables LSP studies in drylands, and highlighted important scale effects driven by within canopy VI variation. With high-resolution imagery the open canopies of drylands are beneficial as they allow for straightforward identification of individual plants, enabling the tracking of phenology at the individual level. Drylands thus have the potential to become an exemplary environment for future LSP research.

## 1 Introduction

Land surface phenology (LSP) enables ecosystem scale tracking of the drivers and consequences of a changing climate. Satellite sensor-derived vegetation indices (VIs) track the progression of green vegetation throughout the year, and from this time series the seasonal transitions between the dormant and growing seasons are derived [1–8]. Numerous studies measure the short- and long-term LSP trends, linking them to drivers such as weather and climate, land cover change, and disturbances [9]. Yet the use of LSP is limited outside regions with adequate vegetation cover and plants with distinct seasonal change. Temperate deciduous forests have the most distinct LSP signal, while in other vegetation types LSP is difficult to discern due to combinations of low signal-to-noise ratio between growing and dormant season VI, high snow cover, and low vegetation cover [10, 11].

In dryland ecosystems low vegetation cover is the primary limitation in detecting land surface phenology [12]. High soil fractional cover can cause the growing season VI to be indistinguishable from the dormant season VI, making phenology extraction impossible. Additionally, some areas with adequate vegetation cover are dominated by evergreen vegetation, thus can have a growing season VI too low to reliably detect seasonal transitions [13]. Even in areas with high vegetation cover and plants with distinct signals, phenology may be impossible to detect some years due to little to no plant productivity from inadequate precipitation. These limitations cause drylands to be excluded in many large scale LSP analyses [14–18], while other analysis include them without any regard for their detectability and resulting bias [19, 20]. Most studies focusing on drylands evaluate aggregate or peak annual VI as opposed to distinct seasonal transitions [13, 21], and even then can occasionally have inconclusive results [22, 23].

Drylands cover 41% of the Earth's surface [24] and reliable measurement of LSP in drylands are needed in understanding these ecosystems and the large-scale roles they play in global processes. Drylands account for much of the long term trends in increased carbon uptake globally [25] and leaf phenology drives much of the inter-annual variation in water and carbon uptake in dryland systems [26, 27]. One quarter of terrestrial vertebrates use drylands to some degree [28] and changing plant phenology can have cascading effects on primary consumers and higher trophic interactions [29]. Additionally, dryland rangelands support 35-84% of livestock globally, thus are a vital source of ecosystem services [30].

Here we explore key attributes of VI time series and how they affect reliable detection of LSP in dryland ecosystems. We use a spectral mixing model to simulate LSP with varying ranges of plant fractional cover, seasonal VI amplitude, and VI uncertainty to determine the feasibility of detecting LSP in different scenarios and with different detection algorithms. We then use high spatial resolution, multi-temporal unmanned aerial vehicle (UAV) imagery to verify some of these relationships. Finally we discuss the feasibility of detecting dryland LSP currently and in the future.

## 2 Methods

### 2.1 Model Simulation

First we describe a conceptual model of an annual dryland VI time series and how it translates into phenology. Using spectral mixing, and assuming a linear aggregation, the VI at a pixel scale for a single date can be decomposed into the following parts:

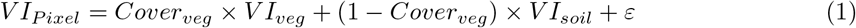

Where *Cover_veg_* is the fractional vegetation cover within the pixel, *VI_veg_* and *VI_soil_* are the average VI values of the endmembers of all plants and soil, respectively, and *ε* is the error term from uncertainty in sensor view angle, shadows, and atmospheric interference [31, 32]. Extraction of phenological metrics relies on the *VI_pixel_* being distinctly higher in the growing season than the dormant season.

Here we explore how variation in the three components in Eq. 1 (i.e., ranges of plant fractional cover, seasonal VI amplitude, and VI uncertainty) affect the ability to detect a distinct phenological signal in sparsely vegetated areas. We start with an idealized annual VI curve derived from a double sigmoid model to represent the VI of single plant canopy (Figure 1, [33]). The curve's amplitude (i.e., the difference between dormant and peak VI values) reflects a combination of plant functional type and leaf area index (LAI). The seasonal amplitude of non-evergreen plants is driven by LAI up to an LAI of approximately 2, while evergreen plants have little to no seasonal amplitude [34, 35]. Higher LAI will generally lead to higher amplitudes, with this effect being more prominent in deciduous plants than evergreen plants [36, 37]. The leaves of senesced grasses have distinctly lower VI than green leaves, thus are expected to behave similarly to deciduous plants [38].

**Figure 1:**
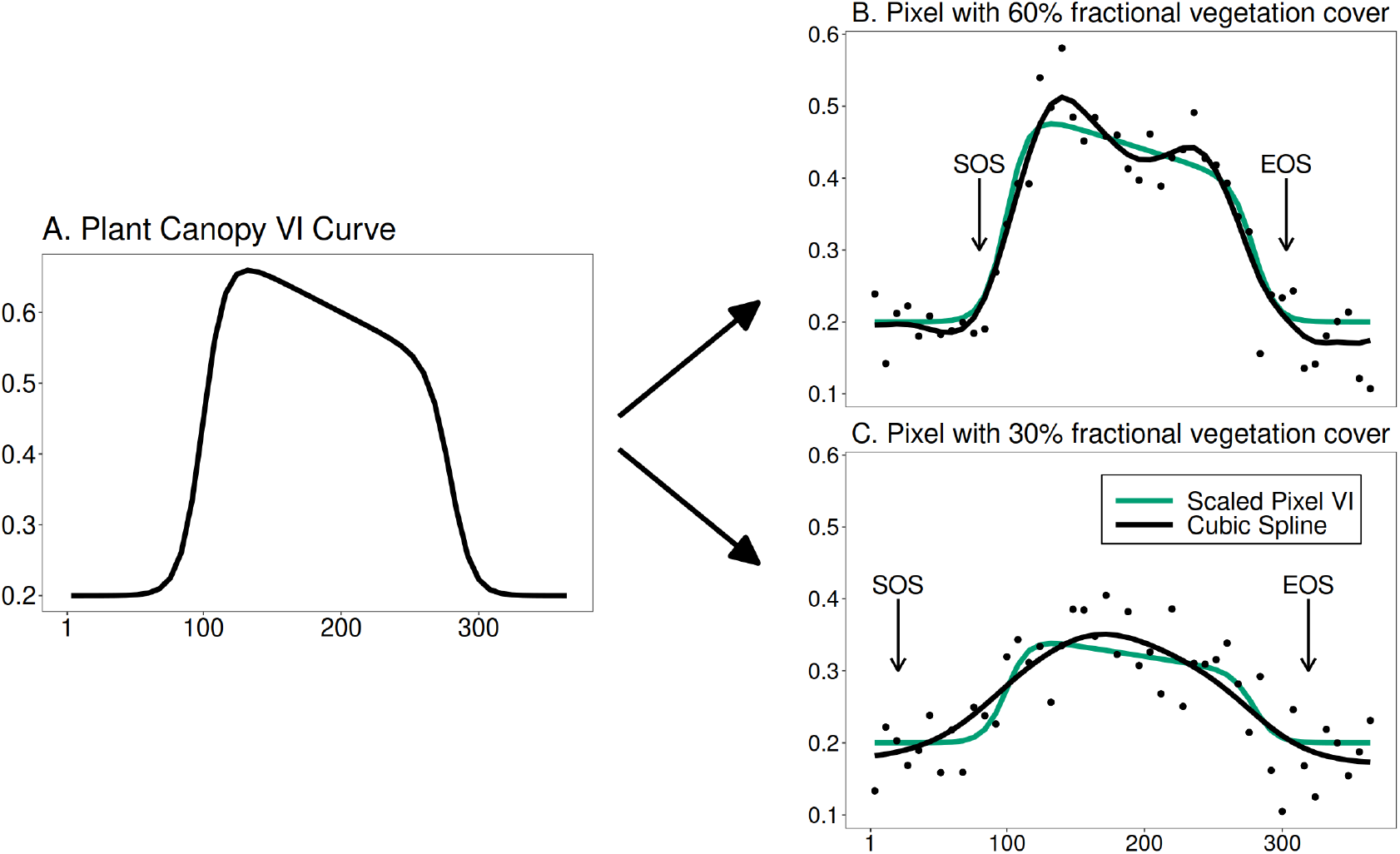
Conceptual diagram of simulating a VI curve with different levels of fractional vegetation cover. Panel A shows an idealized VI curve represented by a plant canopy (ie. 100% fractional vegetation cover). Panels B and C show the same curve scaled to pixels simulating 60% and 30% plant cover, respectively (green lines), and assumes *VI_soil_* remains a constant 0.2. The points represent the scaled curve plus VI uncertainty, with the black line representing cubic spline fit from the points. Phenological metrics (start of season, SOS, and end of season, EOS) can then be extracted from the cubic spline curve. The process is repeated with different amplitudes (ie. the curve height in panel A), plant cover values, and uncertainty size.

Using the curve values at 8-day intervals for a full calendar year, we apply Eq. 1 with fractional vegetation values of 60% and 30%, then include gaussian noise with a mean of 0 and standard deviation (SD) of 0.02 to represent *ε*, resulting in an 8-day VI time series for a full calendar year (Figure 1B, 1C points). A cubic smoothing spline is then fit to the 8-day *VI_pixel_* values and phenology metrics extracted using a threshold of 10% of the relative maximum value. Figure 1 shows how the phenological metrics change due solely to differences in fractional vegetation cover, even though the underlying plant phenology, *VI_veg_*, is the same. In this example there is still adequate amplitude to detect transitions, but at a low enough fractional vegetation cover the V*I_pixel_* amplitude will be too low to detect transition dates.

For the simulation analysis we repeated this process while varying the three different factors from Eq. 1: 1) the fractional vegetation cover (*Cover_veg_*), using values from 0-100%, 2) the amplitude of *VI_veg_* from dormant to peak, adjusted using a parameter in the double sigmoid model, and 3) the size of the error term *ε* using values of 0.01 and 0.04 for the standard deviation of the gaussian noise. For each combination we repeated the process 100 times. A low growing season VI relative to the dormant season is one of the main challenges in dryland LSP [37]. Phenological extraction methods account for this in several ways. A common approach is discarding pixels which do not exceed a threshold of 0.1 VI units between the dormant season and peak growth (i.e., a minimum *VI_pixel_* amplitude of 0.1; [6, 11, 14, 18]), while others have used statistical power methods of the time-series to filter out pixels with low signal-to-noise [39]. Here we evaluate the detectability of LSP in two ways: 1) The frequency at which the *VI_pixel_* meets or exceeds an amplitude of 0.1 VI units, and 2) mean absolute error (MAE) in the estimated phenological metrics start of season (SOS), end of season (SOS), and peak of season (POS).

### 2.2 UAV Study Area, Acquisition, and Processing

We verified the effect of fractional vegetation cover on LSP detection by evaluating LSP of a deciduous shrubland and weekly in situ observations of canopy greenness. We acquired UAV imagery for a long-term study site (NORT) on the Jornada Experimental Range (JER) for eight dates in 2019. The JER is located in the Chihuahuan Desert near Las Cruces, New Mexico, U.S., and is a low-diversity, mixed perennial grassland and evergreen/deciduous shrubland that receives on average 254-mm rain annually with 50% falling between July and October [40]. The site is ideal for testing patterns of LSP detectability and accuracy since it has low to moderate fractional vegetation cover consisting primarily of a single plant species.

The NORT site consists of mostly sandy soil and has approximately 30% cover of honey mesquite (*Prosopis glandulosa*) with occasional four-wing saltbush (*Atriplex canescens*) shrubs growing in some mesquite patches and snakeweed (*Gutierrezia sarothrae*) in the interpatch areas. In addition to UAV imagery we also collected weekly in situ observations of mesquite canopy greenness following USA-National Phenology Network intensity protocols (Denny et al. 2014). Each week during 2019, for five focal mesquite shrubs, we estimated the percentage of green leaves within each individual shrub canopy in seven bins (0%, 1-4%, 5-24%, 25-49%, 50-74%, 74-94%, 95-100%).

To collect aerial imagery we deployed a 3DR SOLO UAV equipped with a Micasense RedEdge camera (https://micasense.com/) and Downwelling Light Sensor (DLS) at an altitude of approximately 28 meters above ground level resulting in a ground sampling distance (GSD) of 1.8 cm. The total image extent was 2.1 ha. Aerial surveys were conducted on eight days in 2019 (April 2, April 25, July 5, August 14, September 26, October 16, November 13, December 20) to characterize mesquite phenology. Flights were planned using Mission Planner software, conducted at mid-day (between 10 a.m. and 12 p.m.); given an east-west corridor pattern, forward lap of 85% and side lap of 75% ensured sufficient pixel overlap between adjacent images for proper alignment and mosaicking. Images of a 6” x 6” Micasense Calibrated Reflectance Panel were taken before each flight for radiometric calibration. We use fixed ground control points within the image footprint as the basis for the geometric correction; ground control points were previously surveyed using a Trimble Geo 7X handheld Global Positioning System. Ortho-mosaics of drone imagery were created using Pix4D software (version 4.2.27, https://www.pix4d.com/) using the following steps:

1. Initial processing: Keypoints Image Scale, full; Matching Image pairs, Aerial Grid or Corridor; Matching Strategy, Use Geometrically Verified Matching; Targeted Number of Keypoints, Automatic; Calibration, Alternative; Rematch, Automatic.
2. Point Cloud and Mesh: N/A
3. DSM, Orthomosaic and Index: Resolution, Automatic; DSM Filters, Use Noise Filtering, Use Surface Smoothing (Sharp); Orthomosaic, GeoTIFF; Radiometric Processing and Calibration, Camera and Sun Irradiance; Resolution, Automatic; Downsampling Method, Gaussian Average; Reflectance Map, GeoTIFF, Merge Tiles.

After initial alignment, GCPs in aerial imagery were manually selected and geolocation tags re-registered with previously surveyed, sub-centimeter geolocation information acquired with the Trimble GPS. The project is then re-optimized with the new geolocation information for more accurate results with RMSE errors ranging from 0.10 to 0.02 meters.

To compute the reflectance value of each pixel, Radiometric Processing and Calibration correction type was set for Camera and Sun Irradiance in orthomosaic and radiometric correction step 3 above. This function in Pix4D calibrates images using irradiance and sun angle measurements obtained from the DLS and the calibration reflectance panel [41]. Reflectance values were used to calculate the normalized difference vegetation index (NDVI) images for the study site for each image date.

Using this time series of NDVI rasters we developed a sampling routine to test the detectability of phenology with respect to fractional vegetation cover similar to the simulation study. We randomly placed 12,000 simulated pixels across the site with sizes of 2, 4, 8, or 16 meters. We then extracted the average NDVI within each one for each image date. This random placement allowed us to produce 12,000 annual NDVI time series representing a range of fractional cover for honey mesquite across several pixel resolutions. The average shrub diameter is approximately 8m, so multiple pixel sizes allow us to test both L- and H-resolution LSP detection [42]. The 16m pixel size represents the L-resolution case in which image elements (i.e., honey mesquite shrubs) are smaller than cell resolution size and cannot reliably be detected individually. Whereas the 2m, 4m, or 8m pixel sizes represent the H-resolution model where honey mesquite elements are larger than cell resolution size and can be reliably resolved in the image. As in the simulation study, each annual time series was fit with a cubic smoothing spline and we then calculated whether the smoothed time series exceeded a 0.1 NDVI threshold and extracted phenology metrics using a 10% of relative max threshold method and the change rate method as described below.

To determine the fractional vegetation cover within each simulated pixel we developed a map of shrub canopy cover for the site using hand annotation of an RGB image, resulting in a raster at the same extent and resolution of the NDVI imagery where each pixel is classified as either mesquite canopy or soil (Figure S1). Fractional vegetation cover was then calculated as the proportion of mesquite pixels within each simulated 2-16 m pixel. To calculate the MAE of the UAV imagery-derived phenology estimates, we used the weekly in-situ observations of mesquite canopy greenness at the site as described above. We fit a single cubic smoothing spline using observations from all five individuals with the midpoint of each percent canopy cover bin. From the spline we calculated the DOY for SOS and EOS using the two methods described below.

### 2.3 Detection Methods and Analysis

Many remote sensing algorithms have been developed in recent decades for estimating transition dates from VI time series, and can be categorized in four broad categories: threshold, curvature and inflection, trend, and priori curve-based approach [8, 43]. Here we use two approaches to evaluate LSP dynamics in drylands: a threshold and inflection (max rate of change) model. The threshold approach uses either fixed or dynamic thresholds of VI values to identify transition dates, is an established and simple methodology, and has been used to produce phenology data products from the Advanced Very High Resolution Radiometer (AVHRR), Moderate Resolution Imaging Spectroradiometer (MODIS), and the Harmonized Landsat and Sentinel-2 (HLS) data [3, 6, 7]. The inflection approach (referred to here as the change rate method) detects the inflection points that show the quickest changes in the VI time series, thus does not depend on a priori thresholds. Curvature and inflection approaches can capture transition dates with relatively small changes in the VI time series, and the curvature method has been used to produce Collection 5 MODIS phenology and VIIRS phenology [1, 44]. Both of these approaches are commonly used in LSP studies [14, 18, 19].

For the threshold method SOS (EOS) was estimated as the DOY when the smoothed VI curve exceeded (fell below) a 10% threshold of the relative maximum VI value [45]. For the change rate method SOS (EOS) was estimated as the DOY when the smoothed VI curve had the highest (lowest) rate of change using the numerically calculated first derivative [46]. The POS estimate was the DOY of the maximum smoothed VI value, thus was independent of the SOS/EOS method. For the MAE of the simulation study the true metrics are derived using the respective method of the original *VI_veg_* annual curve. Since we are using a simulated VI it can be interpreted as either the enhanced vegetation index (EVI) or NDVI. All code for this analysis is available in the Zenodo data repository (https://doi.org/10.5281/zenodo.4777207).

## 3 Results

### 3.1 Model Simulation

When the fractional vegetation cover is 10% or less the *VI_pixel_* curve rarely exceeds the 0.1 threshold between the dormant and peak growing seasons (Figure 2). Even an extremely “bright” plant with a *VI_veg_* amplitude of 0.8 will meet the 0.1 threshold less than half the time. At 60% fractional vegetation cover plants with *VI_veg_* amplitude of 0.2 or higher consistently exceed the needed threshold. Plants with a *VI_veg_* amplitude of 0.1, which might be seen in some evergreen species, never consistently exceed the 0.1 *VI_pixel_* threshold even with nearly 100% fractional cover.

**Figure 2:**
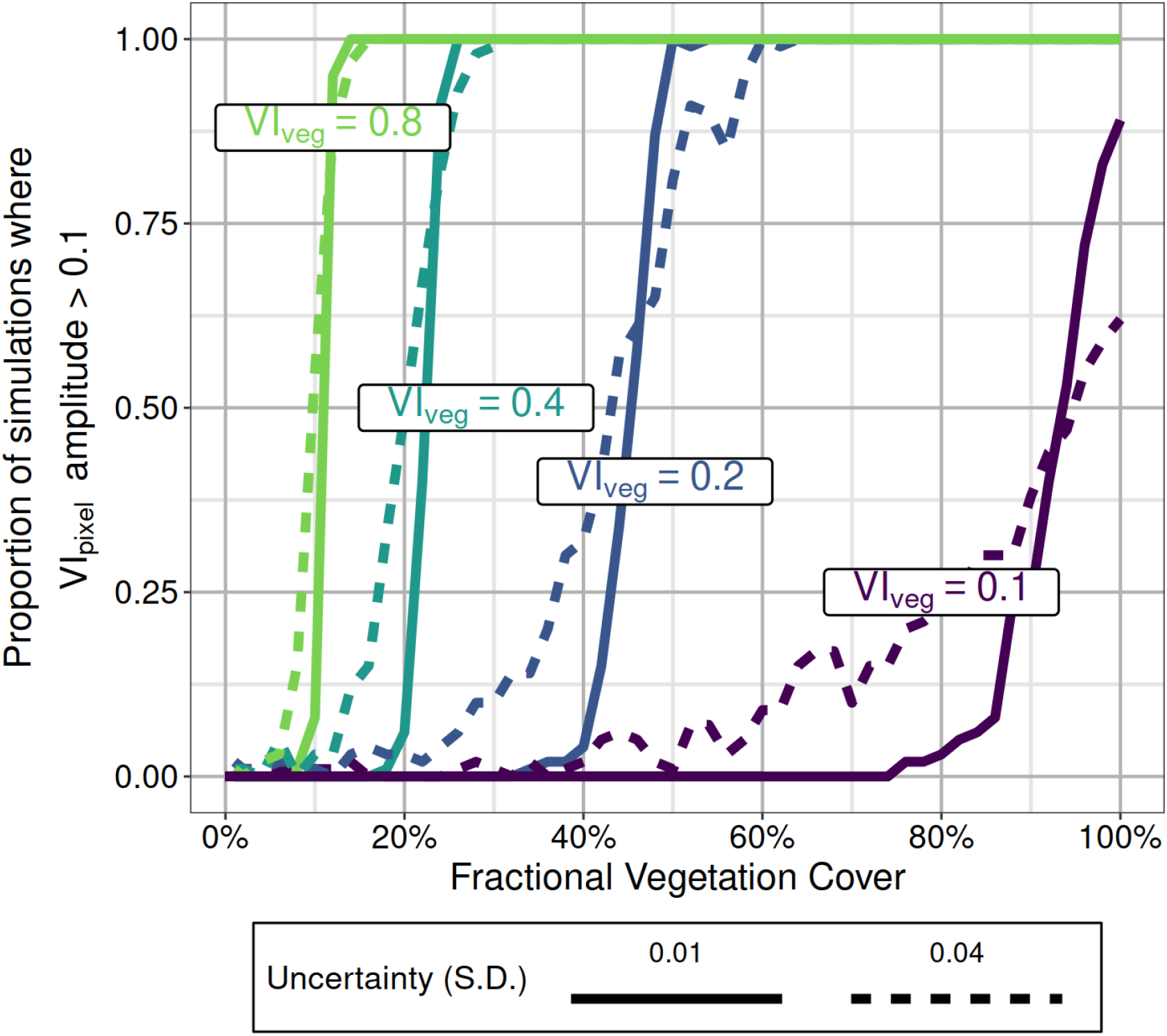
The proportion of simulation runs where the final *VI_pixel_* amplitude exceeds the 0.1 threshold. Colors indicate starting amplitudes of the pure endmember *VI_veg_* curve. Uncertainty is the standard deviation (SD) of the zero centered gaussian noise of the model.

Increasing the uncertainty dampened the effect of increasing fractional vegetation cover. When the uncertainty SD was 0.04, and combined with low fractional vegetation cover, a larger proportion of simulation runs exceeded the *VI_pixel_* 0.1 threshold, with the reverse happening at higher fractional vegetation cover (Figure 2). At low and high fractional vegetation cover VI uncertainty had little to no effect on whether the 0.1 threshold was exceeded. Thus, VI uncertainty propagates into uncertainty in detecting dryland LSP, with this effect being highest at intermediate levels of fractional vegetation cover. For example a pixel with low vegetation cover may, in some years, have seemingly detectable LSP due solely to chance instead of significantly higher productivity.

Even with a phenological pattern that is consistently detectable, the combined effects of soil background, VI uncertainty, and methodology still cause some level of bias in the estimated phenological metrics. For example, given a plant with an *VI_veg_* amplitude of 0.2 and 60% fractional cover, all phenological metrics had mean average error (MAE) of at least 10 (and up to 50) days in our simulations using a 10% of maximum threshold (Figure 3). The MAE of estimates improved with increasing vegetation cover, with increasing *VI_veg_* amplitude, and from using the change rate instead of threshold method for SOS and EOS estimation. The highest errors were seen with low *VI_veg_* amplitude and high (0.04) VI uncertainty. POS estimates were more accurate than SOS or EOS in most instances when using a threshold method, especially when the *VI_veg_* amplitude was low (0.4 or 0.1). The change rate method outperformed the threshold method in all instances except one, when estimating EOS with a *VI_veg_* amplitude of 0.1. The change rate method was particularly accurate in estimating SOS of this simulated data, especially when fractional vegetation cover was low. Increasing the VI uncertainty from 0.01 to 0.04 increased the MAE of estimates in all cases regardless of fractional cover, *VI_veg_* amplitude, or methodology.

**Figure 3:**
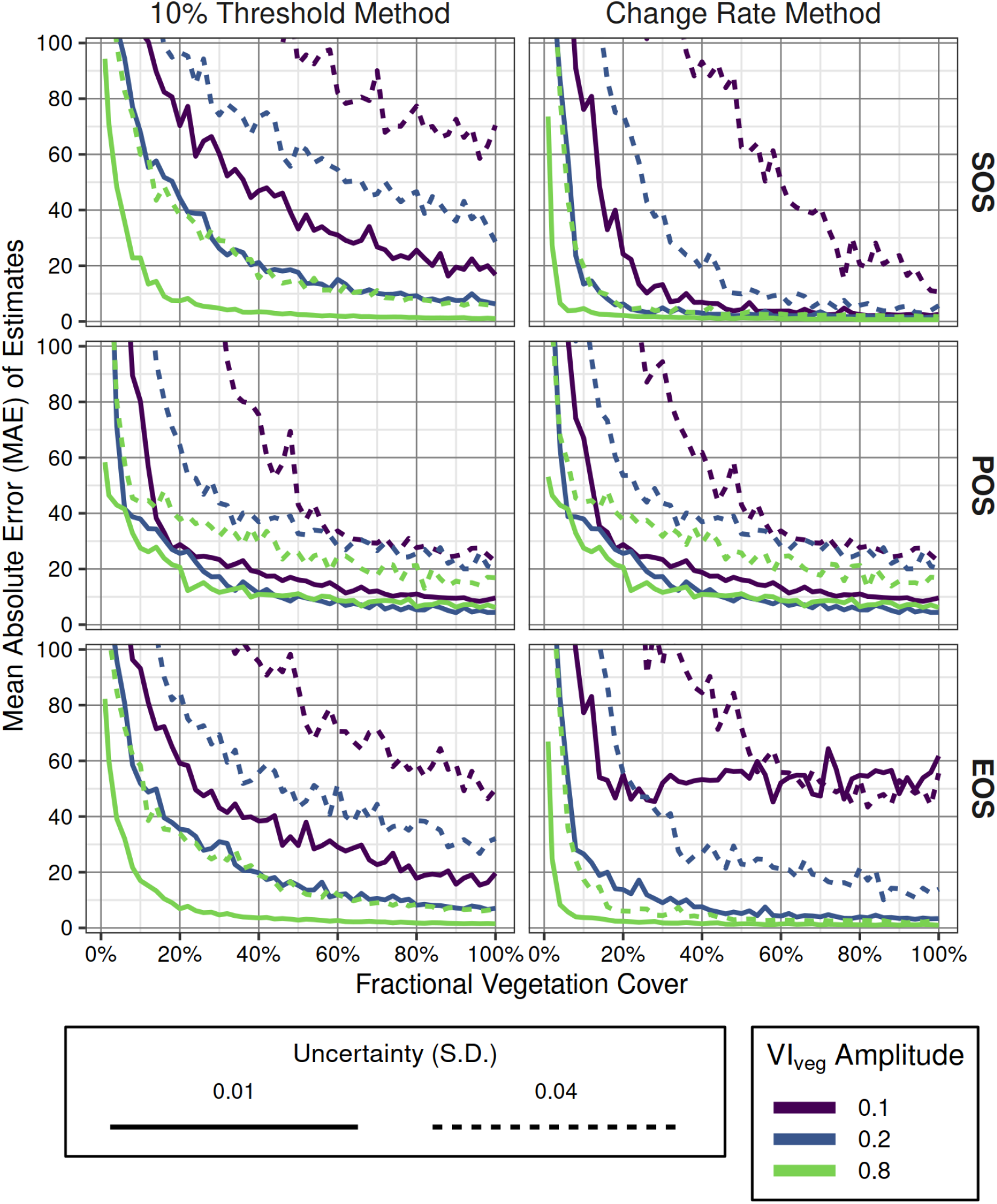
Mean absolute error (MAE) of three phenological metrics estimated from simulated annual vegetation index time series. Start of season (SOS), peak of season (POS), and end of season (EOS). Colored lines indicate the amplitude of the endmember vegetation (*VI_veg_*). Uncertainty is the standard deviation (SD) of the zero-centered gaussian noise of the model. Percent of maximum thresholds are those used in the SOS and EOS estimations. The *VI_veg_* amplitude of 0.4 is excluded for clarity.

### 3.2 UAV Imagery

Annual mesquite NDVI exceeded a 0.1 minimum amplitude consistently when fractional vegetation cover was between 20% and 60%, depending on the simulated pixel size (Figure 4). Larger pixel sizes require less fractional cover to consistently exceed the 0.1 amplitude. This discrepancy with pixel size is similar to what is seen with increasing VI uncertainty in the simulation analysis, but here reflects the high variation in NDVI within the mesquite canopy. For example, given a single mesquite shrub 8m in diameter, 2m pixels randomly located within the canopy can have a wide range of NDVI values for a single acquisition date, resulting in varying amplitude given the same fractional cover (Figure S2, S3). Larger pixel sizes essentially have larger samples to better estimate the mean NDVI of canopies. This leads to a scale dependence for LSP detectability at these extremely fine-scale spatial resolutions. At the 16m pixel size the NDVI variation is reduced enough to result in a relationship between fractional vegetation cover and LSP detectability which most reflects our simulated results.

**Figure 4:**
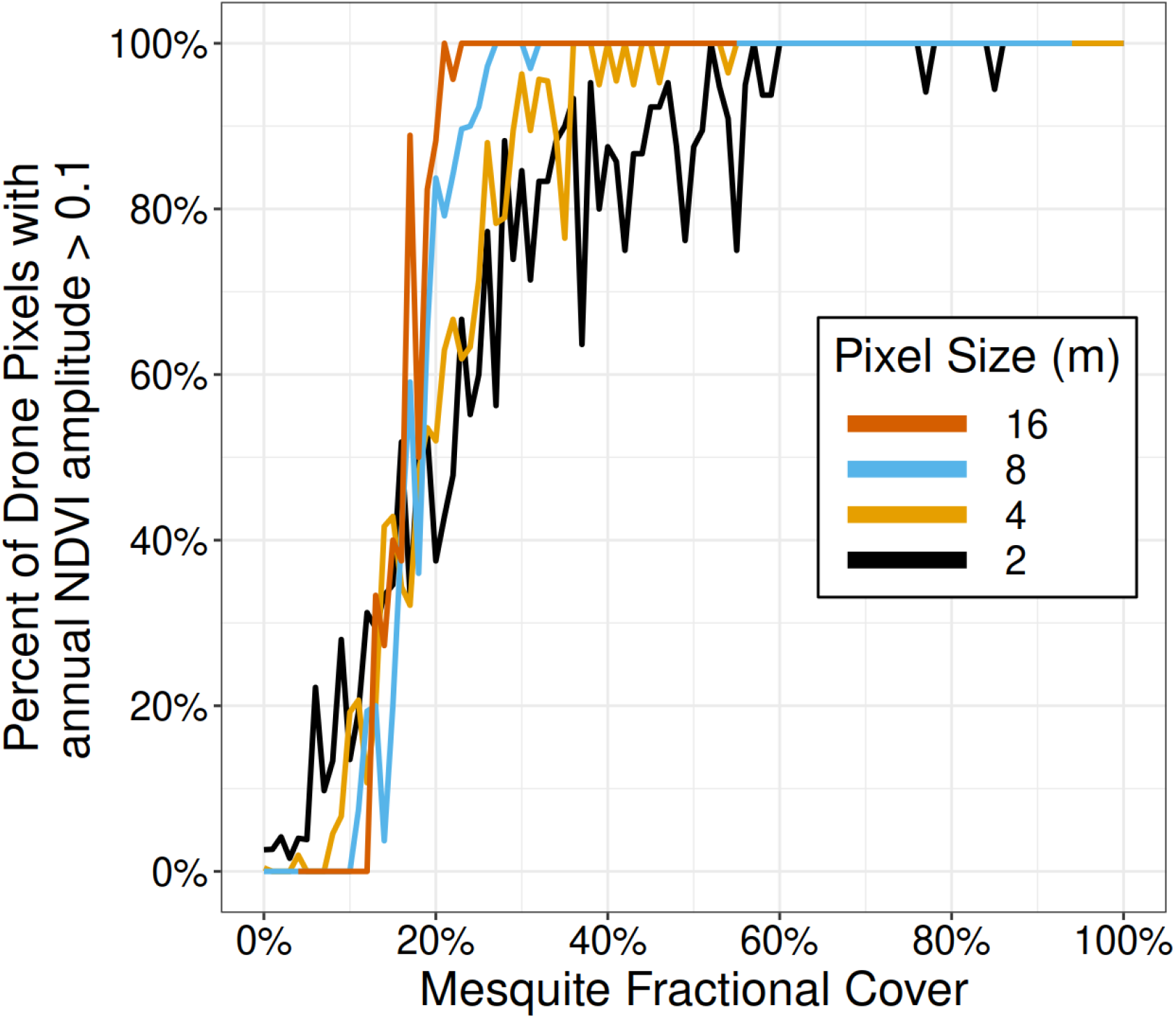
The proportion of pixels where the annual NDVI amplitude exceeds the 0.1 threshold. Colors reflect the simulated pixel size in meters.

As in the simulation study, errors of LSP derived transition dates decreased with increasing vegetation cover for all metrics (Figure 5). Because of the scale effect from canopy variation in NDVI, smaller pixel sizes had large variation in error rates for any given fractional cover. Using pixel sizes of 8 or 16m the MAE stabilized at approximately 20% mesquite cover for SOS and EOS using the threshold method. The change rate method produced slightly more accurate estimates when mesquite fractional cover was less than 20%, but equivalent or less accurate estimates at high mesquite cover. The MAE for POS, which was independent of the threshold or change rate methods, was never below 40 days and commonly greater than 60 regardless of pixel size. This suggests a dissimilarity between POS from UAV imagery versus in-situ observations.

**Figure 5:**
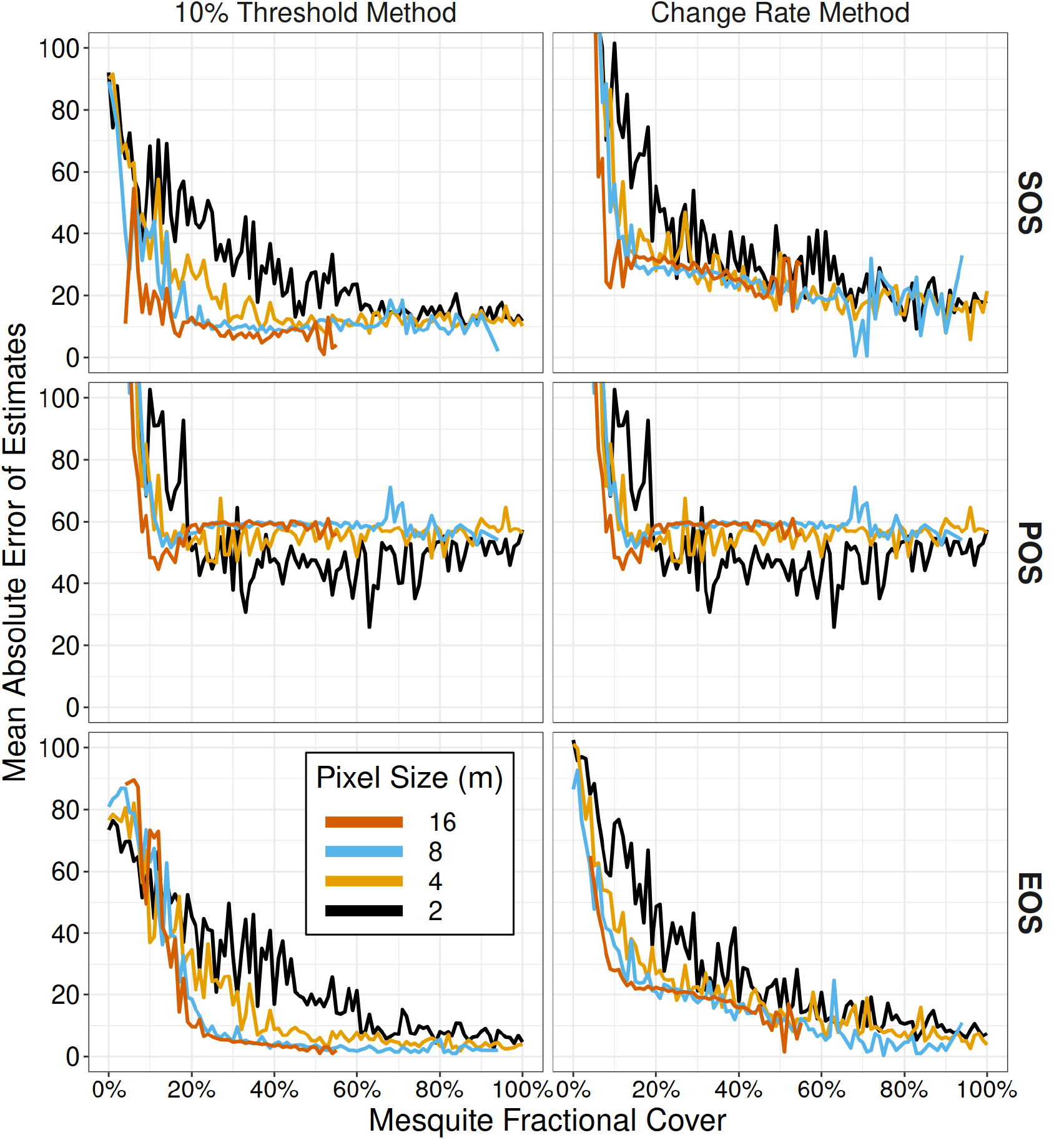
The mean absolute error (MAE) of phenological estimates from 12,000 simulated pixels, with sizes from 2-16m, in an annual UAV imagery time series. Phenological metrics are the day of year for start of season (SOS), end of season (EOS) and peak of season (POS). True metrics were obtained from weekly in-situ observations throughout the year.

## 4 Discussion

With simulated VI curves we showed the limitations in detectability of dryland LSP with respect to fractional vegetation cover, VI amplitude, and VI uncertainty. The required fractional vegetation cover to detect phenology depends on the underlying plant traits, with evergreen plants needing higher cover than non-evergreen plants. For example our UAV analysis showed that with 8m or 16m pixels 20-30% deciduous shrub cover is needed to consistently detect LSP, while Peng et al. 2021 [47] found that approximately 50% cover is needed for consistent detection in evergreen dominated shrublands. Even when the seasonal amplitude of a pixel exceeds a minimum amplitude threshold, phenological metrics of sparsely vegetated drylands can still have considerable error due to other factors. Our UAV imagery analysis validated these results, and also highlighted how within canopy VI variation affects LSP detection in high-resolution imagery. The algorithm used for transition date estimation affected results as well, with the change rate method outperforming the threshold method with simulated data, and the threshold method having better or equivalent results than the change rate method with UAV imagery time series.

The three primary factors affecting LSP detection and accuracy vary across ecosystems, sensors, and with plant community composition. They also interact such that the detectability is not uniform across vegetation types with similar characteristics. For example 40% fractional cover may be adequate to detect LSP in a deciduous shrubland, but likely not in an evergreen shrubland. Neither may have detectable LSP with some satellite sensors due to higher VI uncertainty. The methodology and algorithms used can also affect results, and some methods may be more suitable depending on the underlying time series characteristics. Here we explore these factors in detail, how they vary, and how they interact.

### 4.1 On fractional vegetation cover

Low fractional vegetation cover is a primary limitation in detecting LSP in drylands [37]. There is high spatial and temporal variability in vegetation cover throughout arid regions. To highlight where vegetation cover is the limiting factor in LSP detection we calculated the average and standard deviation of total fractional vegetation cover in drylands in the western United States across 20 years (Figures 6 & 7). There is a general gradient of increasing fractional cover moving from south to north in deserts of the USA. The Sonoran and Mojave desert ecoregions have the lowest fractional cover, with median values of approximately 20%. In the northern regions there are large areas exceeding 60% fractional cover. Whether this translates to detectable phenology depends on the plant composition and resulting *VI_veg_* amplitudes. Even at 100% fractional cover there may be no LSP signal due to, for example, low *VI_veg_* amplitude in evergreen plants (Figure 2). These dynamics can result in confounding trends where there may be higher LSP detectability in areas with lower fractional vegetation cover.

**Figure 6:**
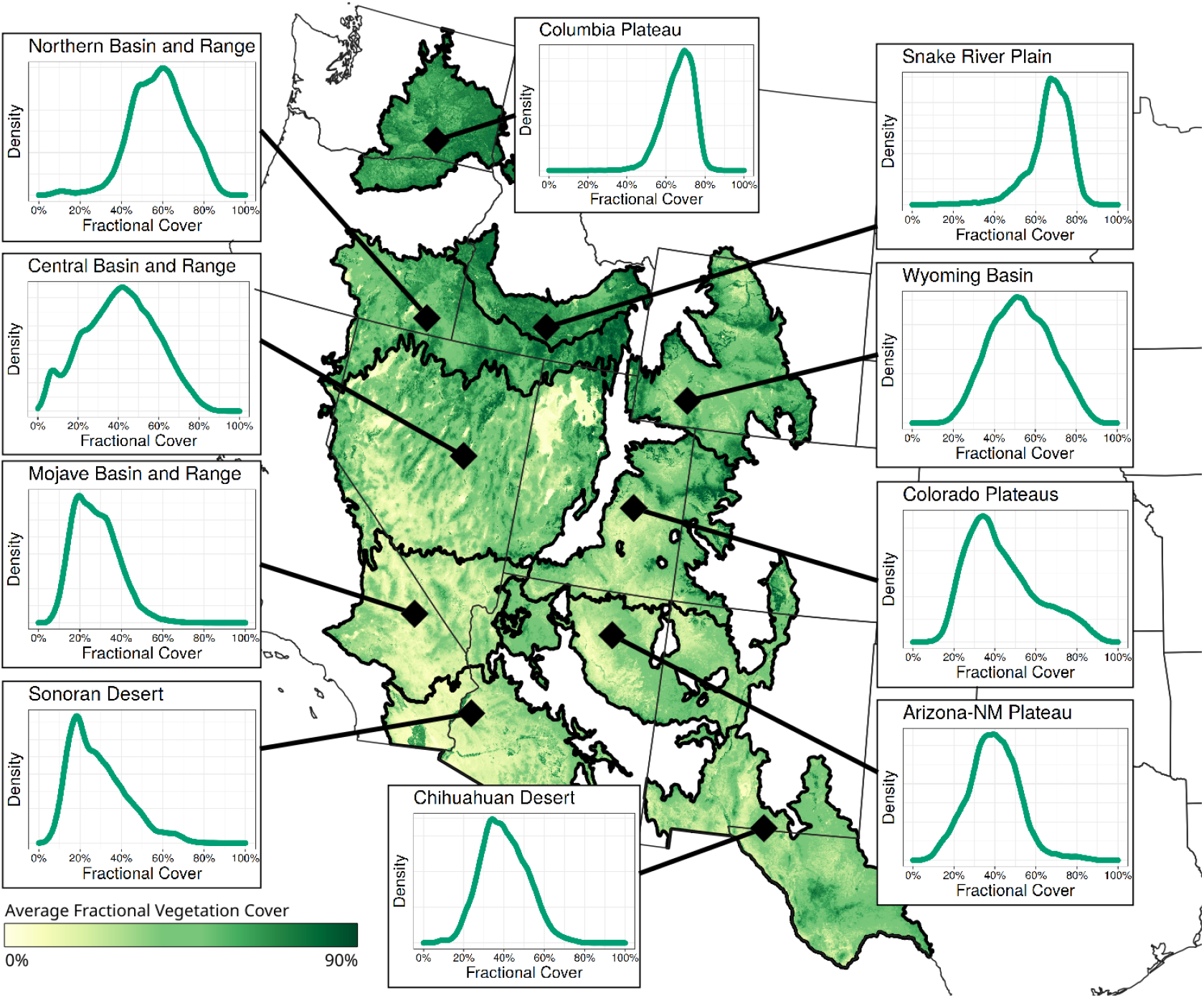
Average annual fractional vegetation cover across Level 3 North American Desert Ecoregions in the United States. Cover represents the average cover from 2000-2019, where the annual cover is the sum of trees, shrubs, annual and perennial forbs and grasses. Inset figures are density plots of the average annual fractional cover for the respective ecoregion. Derived from the dataset in Allred et al. 2021 [92].

**Figure 7:**
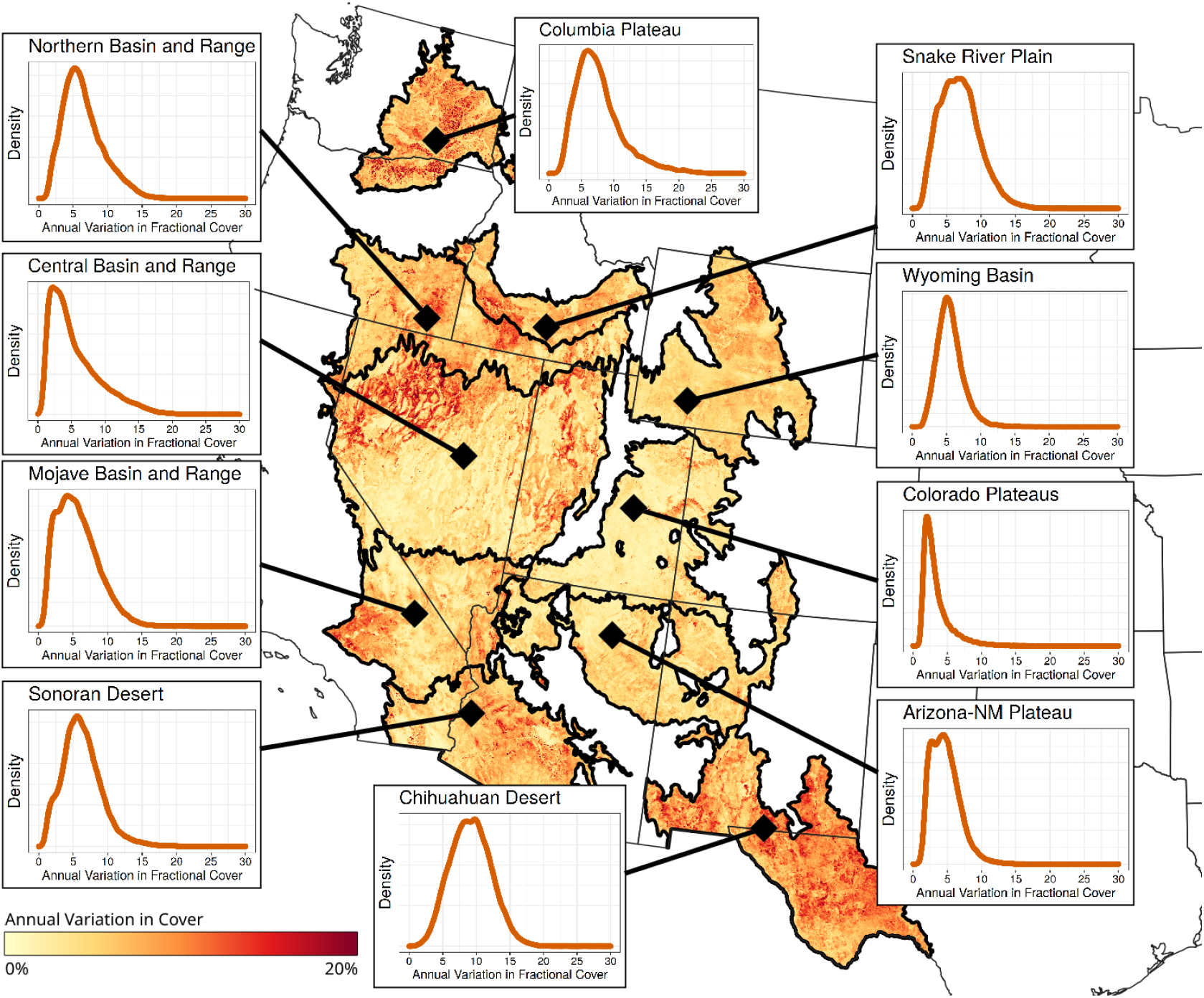
Variation in annual fractional vegetation cover across Level 3 North American Desert Ecoregions in the United States. Cover represents the standard deviation in cover from 2000-2019, where yearly cover is the sum of trees, shrubs, annual and perennial forbs and grasses. Inset figures are density plots of the year to year standard deviation in fractional vegetation cover for the respective ecoregion. Derived from the dataset in Allred et al. 2021 [92].

Year to year variation in vegetation cover is highest in the Chihuahuan Desert ecoregion and portions of the Sonoran and Mojave deserts (Figure 7). Here there is high variability in the germination of annual plants as well as greenup of perennial grasses, both of which are driven by precipitation [48, 49]. Confounding this further is different timings of greenup among plant functional groups in response to precipitation, resulting in two distinct growing seasons [50–52]. The dynamics of this bi-model seasonality was not explored here, but can be accounted for with the correct methods [6, 19].

In the northern ecoregions there are large areas of high variability which can be attributed to agriculture [53]. Outside agricultural areas variation in annual cover in the northern ecoregions is likely driven by disturbance and subsequent changes in plant composition. Large swaths of the Snake River Plain and Central and Northern Basin and Range have high variability due to high severity fires and subsequent annual grass dominance [54, 55]. Here, the detectability of LSP may increase when annual grasses replace evergreen shrubs, since the latter have a lower seasonal amplitude [47].

Changing plant composition can lead to variability in cover and subsequent LSP detectability [56]. From the trends seen across desert ecoregions, changing composition must be considered on two time scales. The first is long-term changes due to disturbance and succession, as seen in northern ecoregions. The second is within year variation where the apparent fractional cover and greenness as seen by satellite sensors is a function of which plants have responded to abiotic drivers [57].

### 4.2 On seasonal amplitude expectations

The seasonal amplitude of plants, the difference in VI between dormancy and peak, drives LSP dynamics. A larger amplitude means that less fractional cover is needed to accurately detect phenological transitions. At the scale of an individual plant seasonal VI amplitude is driven by functional type and leaf area index (LAI).

Canopies of non-evergreen plants, such as grasses, forbs, and deciduous shrubs can have high seasonal amplitudes, since in the dormant season these canopies consist primarily of background soil, litter, and/or senesced vegetation. A higher canopy LAI can increase the seasonal amplitude further, up to an approximate LAI of 2, beyond which VI's tend to saturate [37]. This has implications for long term trends in LSP related to plant growth. Once a plant matures to the canopy LAI saturation point, only further horizontal growth can increase pixel level VI since vertical growth does not decrease overall soil cover [34, 35].

Evergreen plants will have relatively low seasonal amplitude, and thus require significant fractional cover to detect phenology. Assuming LAI remains relatively constant throughout the year then variation in the VI of evergreen plants depends solely on the structure and turnover of leaves, resulting in little to no seasonal amplitude. Indeed, studies of several evergreen shrubs found them to have seasonal amplitudes of less than 0.1 NDVI units [34, 58]. Thus even when fractional cover approaches 100%, pixels with predominantly evergreen vegetation cannot have detectable phenology. Furthermore, pixels with high amounts of evergreen vegetation may occasionally have LSP detections solely by chance (ie. false positives) due to the inherent uncertainty in satellite imagery. Our simulation results show that the resulting phenological metrics in these scenarios, where the VI_pixel_ amplitude rarely exceeds a minimum threshold, can have errors of several weeks or more.

### 4.3 On VI uncertainty

VI uncertainty from shadows, view angle, and atmospheric interference can increase the false positive and false negative rates in LSP detections and increase errors in resulting transition dates [12]. False negatives can occur when a VI time series does not meet the minimum threshold when it otherwise would with zero uncertainty. False positives can occur when a VI time series erroneously exceeds a minimum threshold when it normally would not have with zero uncertainty. Our simulations show that increasing VI uncertainty increases false positives at low fractional cover, and increases false negatives at higher cover (Figure 3). Higher uncertainty also increases error in transition date estimates [59]. Our simulation results showed that increasing VI uncertainty increased the mean absolute error in all scenarios and metrics (Figure 4).

At intermediate levels of fractional vegetation cover VI uncertainty determines LSP detectability, whereas very high or very low fractional vegetation cover leads to a signal to noise ratio which makes VI uncertainty negligible. The upper threshold of fractional vegetation cover at which VI uncertainty becomes negligible for LSP detection depends on the seasonal VI amplitude of vegetation. Vegetation with high seasonal amplitude, such as deciduous plants with high LAI, have a lower threshold of fractional vegetation cover, above which VI uncertainty is negligible for LSP detection. LSP detection of vegetation with low seasonal amplitude, such as evergreen plants, is affected by VI uncertainty even at 100% fractional cover (Figure 2). Indeed, the combination of high VI uncertainty and evergreen dominated landscapes can lead to improbable winter peaks in greenness [60].

In our simulation analysis we characterized VI uncertainty as zero-centered gaussian noise with a constant standard deviation of either 0.01 or 0.04. In reality the standard deviation of VI uncertainty can range from less than 0.01 to over 0.1 depending on the specific index used, its magnitude, and the sensor [61–63]. VI uncertainty can also vary between the dormant and growing season due to changing VI magnitude. Uncertainty decreases with increasing VI magnitude for NDVI and most other indices, while EVI uncertainty increases with higher EVI magnitude [61]. Differences in design, degradation, orbital drift, and spatial and spectral resolution lead to different VI uncertainties across sensors [61, 64]. Coarser spatial resolution decreases uncertainty [65], though this does not imply upscaling will produce more precise phenological estimates [8, 66, 67]. Data products which are corrected to surface reflectance have significantly lower uncertainty than top of atmosphere reflectance, though the uncertainty is still high enough to produce the patterns seen here [62].

### 4.4 On methodology

The algorithms used for LSP detection in drylands need further study, ideally to identify the underlying mechanism which makes one method more suitable than others. Using simulated, and idealized, data we found the change rate method generally outperformed the threshold method except for EOS estimates of low *VI_veg_* amplitudes. This was likely the result of the gradual greendown in underlying double sigmoid used to generate the simulated data, where after adding VI uncertainty the steepest part of which was highly variable in the resulting cubic spline smoother. With UAV imagery the threshold method produced lower MAE than the change rate method in most instances. This could be due to the low temporal density of only eight UAV flight dates throughout the year. With limited sample size the date of maximum change in the smoothed VI curve can be more variable than the date when a relative threshold is reached. Thus, any evaluation of algorithms for dryland LSP detection must also consider the temporal scale of the sensor in addition to the vegetation and VI attributes. The most suitable algorithm for UAV imagery in a particular ecosystem will not necessarily be the most suitable for satellite based imagery in the same ecosystem.

Other algorithmic improvements may also aid dryland LSP studies. For example, methods which reduce VI uncertainty can be beneficial, since here we have shown this can decrease false positive and false negative detections and also increase the accuracy of the resulting transition date estimates. Propagation of VI uncertainty cannot improve LSP metrics directly, but can provide better context for transition date estimates through confidence intervals, which are rarely used in LSP trend studies [68]. VI uncertainty propagation is difficult to implement though [63], and would likely best be done as additions to Level 2 or higher data products.

### 4.5 On mixed pixels

The single vegetation type used in our simulation and UAV imagery analysis is likely rare in most drylands [69]. Mixed vegetation pixels complicate LSP in several ways beyond what we evaluated here. For example Chen et al. 2018 [56] found that, in a pixel with two vegetation types, the estimated green up date can vary solely due to changing species composition and not changing plant phenology. Dryland plant dynamics complicate LSP detection further since different functional types respond differently to precipitation pulses [57]. For example grasses may green up sporadically, or even not at all, due to the amount and timing of precipitation. Conversely surrounding shrubs, which access deeper water pools, can leaf out consistently every year [70–72]. Combined with low fractional cover this makes sparse vegetation drylands one of the hardest ecosystems for detecting LSP.

### 4.6 On low amplitude errors

Here we have highlighted a little studied aspect of LSP research. As *VI_pixel_* amplitude decreases, either from lower fractional vegetation cover or lower *VI_veg_* amplitude, potential error in transition date estimates increase. This is associated with the inherent parameter uncertainty in smoothing algorithms, leading to a statistical limitation in estimating transition dates. We illustrated this effect in Figure 8, where three VI curves have the same VI uncertainty (SD = 0.01) but different *VI_pixel_* amplitudes. As amplitude decreases, the potential range of onset dates increases. This happens with both a threshold and change rate method, though the change rate method is less sensitive. As seen in both the simulation and UAV imagery analysis this leads to higher error potential at lower *VI_pixel_* amplitudes. Confidence intervals, and errors of resulting transition date estimates, can be decreased with different temporal composite methods and/or with smaller temporal resolutions [73]. A best practice would also be to incorporate transition date confidence intervals into LSP studies for more appropriate statistical tests.

**Figure 8:**
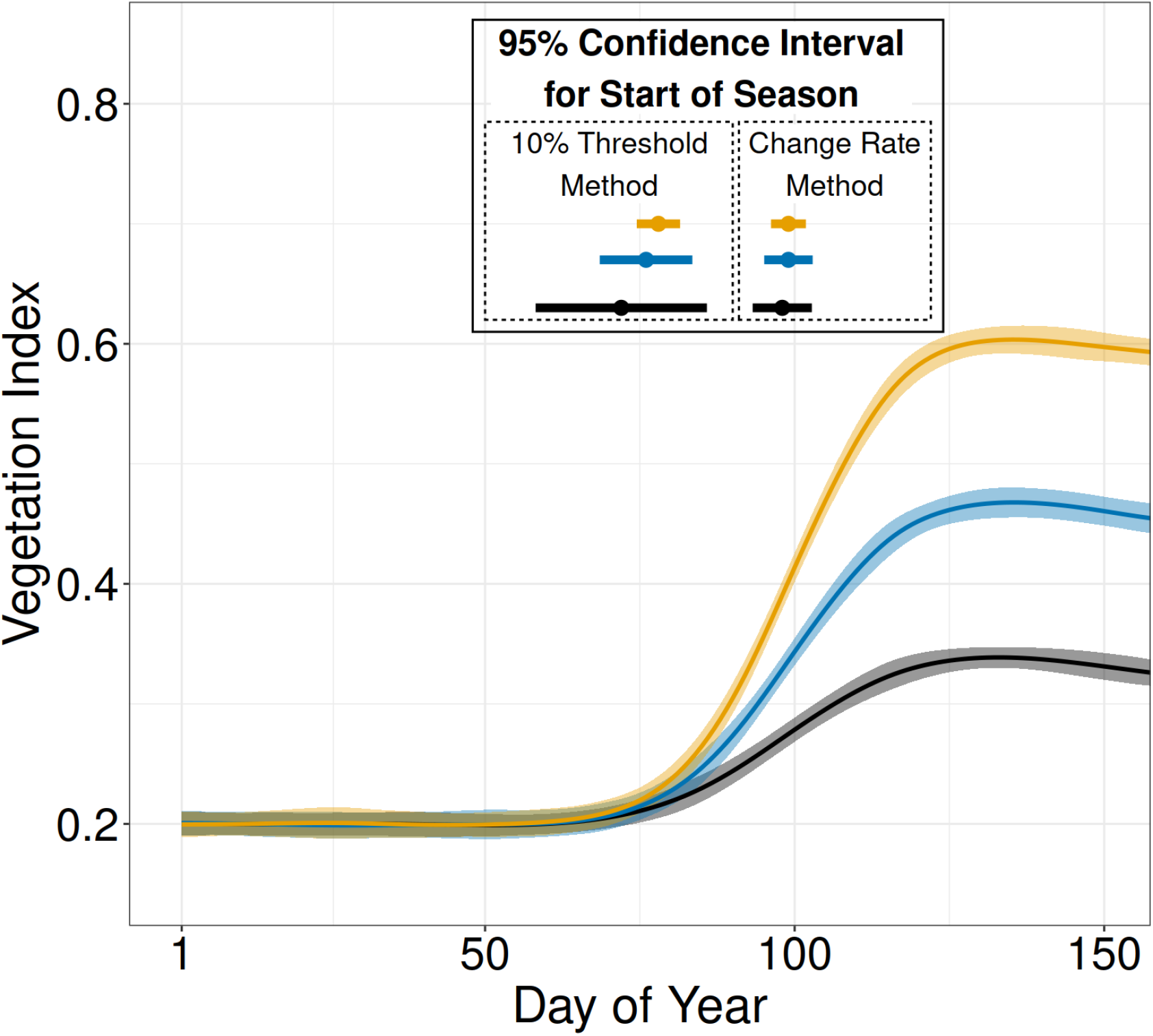
Three simulated VI curves with the same VI uncertainty (SD = 0.01) but different seasonal amplitude. Ribbons represent the 95% confidence interval around each VI curve as estimated from a cubic smoothing spline. Start of season estimates align with the x-axis, and are based on either a 10% of maximum threshold or the change rate method. Horizontal lines for the start of season estimates represent the mean and 95% confidence interval corresponding to the respective method and same colored VI curve, and align with the x-axis.

### 4.7 Moving Forward

Due to low seasonal amplitude the timing of seasonal transitions cannot be accurately estimated in many dryland areas with the current suite of satellite sensors and methodologies, though there is opportunity in some instances. Opportunities exist in areas with high spatial variation in vegetation type and cover, where small patches of vegetation meet or exceed the needed requirements of amplitude and cover. In these areas finer resolution sensors (eg. Landsat & Sentinel-2) can potentially detect LSP where coarser resolution sensors (eg. MODIS, VIIRS, & AVHRR) cannot [47]. Another opportunity is infrequent “super blooms” which occur in several desert regions worldwide and have distinct VI signals [74]. High fractional cover of flowers require special consideration since they have lower VI values than green leaves, but still significantly higher than bare soil [75]. This is also an opportunity to upscale vegetation indices developed for flowers in agricultural areas [76, 77].

In areas where *VI_veg_* amplitude is extremely low the timing of POS is likely the most suitable metric. Here we found estimates of POS showed less bias than those for the SOS or EOS, especially when *VI_veg_* amplitude was low. Other studies have shown the timing and amplitude of POS to be reliable in studying evergreen phenology [13, 78] and annual vegetation biomass [79]. Though caution must still be observed here since low *VI_veg_* amplitude can lead to improbable winter peaks [60].

With high-resolution satellite or UAV sensors sparse vegetation can be advantageous in tracking phenology. Areas of open canopy, which are common in drylands, are the easiest environments for automatic identification of individual plants in high-resolution imagery [80]. By combining high-resolution imagery with frequent overflights the phenology of individual plants can then be tracked. As opposed to landscape level variation in phenology of moderate resolution sensors, high-resolution sensors could thus be used to study phenology within and among species or across different microhabitats in drylands (e.g., [49]). The within canopy variation in VI, usually negligible at coarse resolutions, can affect LSP detectability. Though results from our UAV imagery analysis suggest that aggregating over all pixels within a plant canopy is likely adequate.

Similarly, UAV imagery can also be used to track phenology at the individual plant level. We found a potential shortcoming in using UAV imagery for LSP analysis, where the POS had large errors relative to SOS and EOS estimates. This could be due to the true date of POS being between our flight dates. Another possible cause is variation in timing of leaf maturity and senescence within the plant canopy, causing differences due to the oblique angle or categorical ocular estimates of insitu observations compared to UAV imagery. Understory species within the mesquite canopy may drive an earlier peak when measured by UAV imagery. Thus when using UAV imagery to estimate phenology, one should maximize the number of sample dates, especially since POS can only be estimated retrospectively. As high-resolution imagery becomes more accessible and commonly used, the relationship between fine scale canopy attributes and how they relate to in-situ observations should be explored more [81, 82].

Other data products which are not proxies of greenness show promise in detecting dryland dynamics, since greenness is not always correlated with physiological activity [83]. Solar-Induced Chlorophyll Fluorescence (SIF) is theoretically not affected by high soil cover since soil is non-fluorescing [84, 85]. In practice SIF-derived LSP transition dates, when compared with in-situ observations, have higher errors in areas with low vegetation cover, likely due to the low seasonal amplitude of the resulting annual time series [86]. Vegetation optical depth, derived from passive microwave sensors, is more sensitive to woody and herbaceous foliage than NDVI and can potentially help discriminate between them [23, 87]. Weather and climate data such as air and soil temperature and precipitation would likely be beneficial in constraining transition dates to biologically realistic ranges [60, 88, 89]. Integrating data from multiple sensor types, thereby retaining the relative advantages of each, shows promise in improving the detectability and accuracy of dryland LSP [23, 87, 90, 91].

The definition of LSP detectability, here whether a *VI_pixel_* amplitude exceeds 0.1 VI units, can potentially be improved. The 0.1 threshold is used in numerous studies, yet its origin and suitability is unclear. Here we showed that it indeed excludes instances where errors can be extremely high, yet does not fully minimize errors in all scenarios. A better methodology or framework for LSP detection could help in dryland LSP studies. For example, instead of whether a pixel amplitude exceeds an arbitrary threshold, detectability might be better defined as whether sensors could discern if plants actually exited dormancy. A false negative detection would then occur if plants within a pixel were deemed to have remained in dormancy, and no transition dates were estimated, when in fact there was plant growth and/or productivity. Theoretically this would make detectability invariant to fractional vegetation cover. Re-framing the detectability of LSP like this, or in other substantial ways, can lead to more innovative research for studying dryland LSP.

## 5 Conclusion

Drylands constitute approximately 41% of the Earth's surface and account for a large amount of ecosystem services and biodiversity [28, 30]. Thus, it is important to understand the drivers and limitations of reliable LSP detection in drylands. As in other studies we found that the interaction of fractional vegetation cover and seasonal VI amplitude of plant canopies are the primary drivers of dryland LSP detectability and accuracy [37, 47]. High vegetation cover leads to higher detectability, except when plants have low VI amplitudes, for example in evergreen vegetation. VI uncertainty associated with shadows, atmospheric interference, and view angle is largely negligible in determining if a VI signal will exceed a minimum amplitude when fractional vegetation cover is either very high or very low. At intermediate levels of fractional vegetation cover VI uncertainty can increase the false positives and negatives of the true VI signal exceeding a minimum amplitude. Regardless of whether a VI signal exceeds a minimum threshold, reducing VI uncertainty can reduce the errors of resulting LSP metrics in all cases. Of the two algorithms used here for LSP estimates, neither performed the best in all scenarios. Studies evaluating the numerous LSP methodologies with a focus on drylands would be highly beneficial, especially if they could identify underlying factors which determine which method performs best in a particular plant community.

High-resolution sensors with sub-meter resolution are a promising path forward for measuring LSP in drylands. These sensors overcome the problem imposed by low vegetation cover, allowing for LSP detection of individual plants. Frequent visits are still needed to adequately capture all seasonal dynamics, which may be problematic when using labor intensive UAVs. High-resolution satellite based sensors will be highly suitable for LSP studies in drylands due to frequent visits and low cloud cover in dryland environments. Since high-resolution imagery also allows for the identification of individual plants, it will be possible to measure LSP at the individual level. Drylands thus have the potential to become an exemplary environment for future LSP research.

## Supporting information

Supplemental Images

## Acknowledgments

This research was a contribution from the Long-Term Agroecosystem Research (LTAR) network. LTAR is supported by the United States Department of Agriculture. DMB and RAB were supported by CRIS #3050-11210-009-00D. We acknowledge the Jornada Basin Long-Term Ecological Research (LTER) site for sustaining the long-term research location (DEB 20-25166). The authors acknowledge the USDA Agricultural Research Service (ARS) Big Data Initiative and SCINet high performance computing resources (https://scinet.usda.gov) and funding from the Scientific Computing Initiative (SCINet) Postdoctoral Fellow program to support SDT. Any use of trade, firm, or product names is for descriptive purposes only and does not imply endorsement by the U.S. Government. USDA is an equal opportunity provider and employer.

## Data Availability

All simulated VI data, simulated pixel time series from UAV imagery, and code for reproducing this analysis, is available in the Zenodo data repository (https://doi.org/10.5281/zenodo.4777207).

## Author Contributions

S.D. Taylor performed the analysis and wrote the manuscript with guidance from D.M. Browning. R. Baca collected and processed the UAV imagery and phenology data. D.M. Browning and F. Gao provided important comments and revised the manuscript.

