## Supplemental Images for "Constraints and Opportunities for Detecting Land Surface Phenology in Drylands"

**Supplemental Images S1-S3**

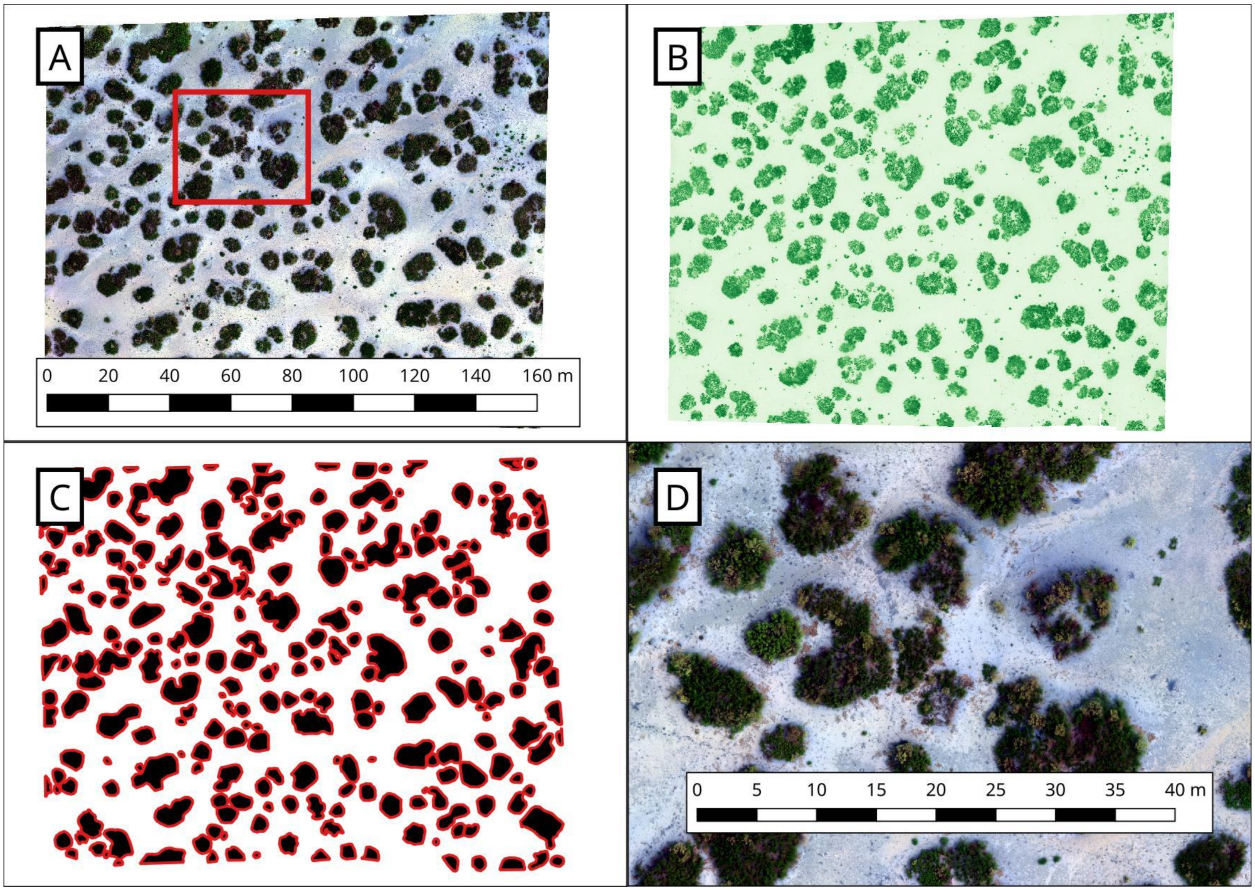


Figure S1: UAV imagery from July 5, 2019 at the Jornada Experimental Range NORT site used in this study. A: The true color RGB image. B: The NDVI image. C: The hand annotated mesquite cover map. D: A zoomed in portion of A represented by the red outline. The scale bar in A is also representative for B and C.

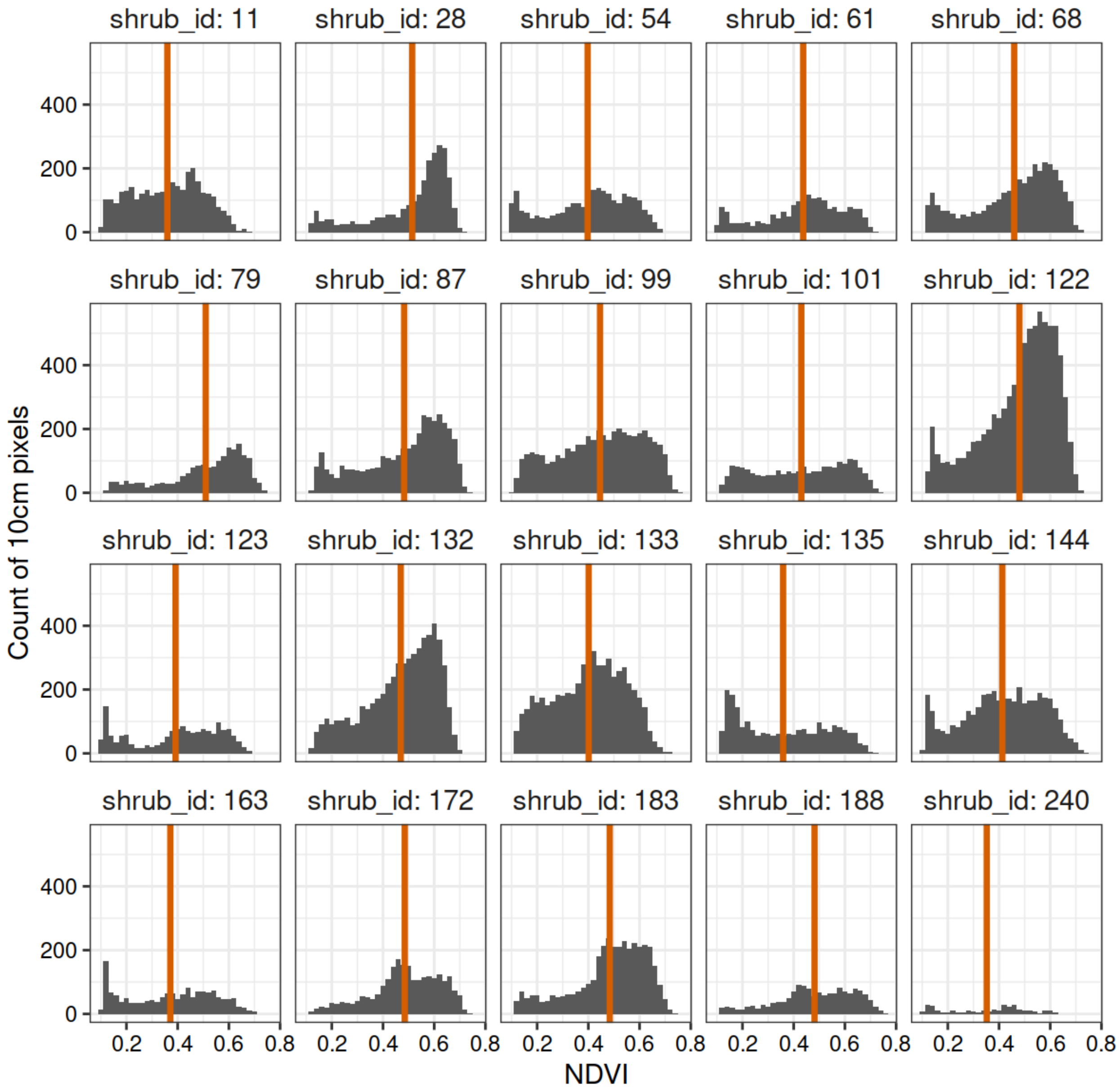


Figure S2: NDVI variation within mesquite shrub canopies. We used outlines of 20 randomly selected shrubs at the NORT site and extracted 10cm NDVI values for a single date representing peak NDVI in 2019. (2019-07-05). Histograms represent the range of NDVI values within the respective shrub canopy, red vertical lines represent the mean NDVI of the entire canopy.

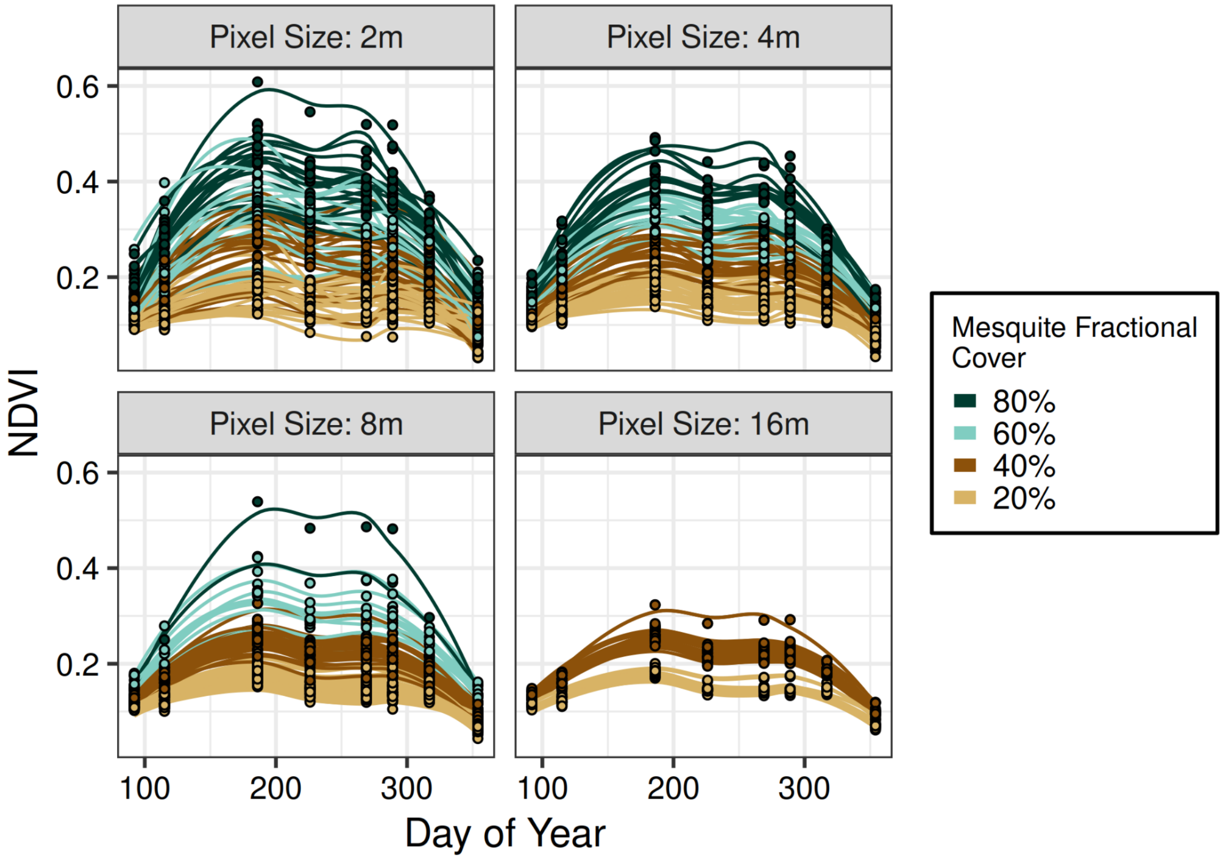


Figure S3: Example NDVI curves from UAV imagery representing 4 pixel sizes (in meters) and 4 levels of fractional vegetation cover.
